# High prevalence of KPC-3 in carbapenem-resistant *Pseudomonas aeruginosa* across multiple clonal lineages in China

**DOI:** 10.64898/2026.08.11.744309

**Authors:** Shanshan Wang, Meilan Li, Zhixuan Chen, Lan Chen, Xingbei Weng, Liang Chen, Bingjie Wang

**Author notes:** These authors contributed equally to this work and share first authorship (Shanshan Wang, Meilan Li). Corresponding author at: Department of Clinical Laboratory Medicine, Shanghai Pulmonary Hospital, Tongji University School of Medicine, Shanghai, 200433, China. **E-mail address** (Xingbei Weng); (Liang Chen); (Bingjie Wang).

## Abstract

**Background:** The epidemiology of *Klebsiella pneumoniae* carbapenemase (KPC)-producing *Pseudomonas aeruginosa* is rapidly evolving in China. While *bla*_KPC-2_ remains the predominant KPC variant in *P. aeruginosa*, *bla*_KPC-3_ has rarely been documented in this pathogen. This study investigated the molecular epidemiology, resistance and virulence characteristics, and plasmid features of *bla*_KPC-3_-producing CRPA isolates collected from a tertiary hospital in eastern China.

**Methods:** A total of 65 non-duplicate CRPA isolates collected in 2023 were subjected to whole-genome sequencing. Antimicrobial susceptibility testing, phylogenetic analysis, plasmid characterization, conjugation experiments, and virulence assays were performed.

**Results:** Among the 65 CRPA isolates, 37 (56.9%) carried *bla*_KPC-3_. These *bla*_KPC-3_-positive isolates belonged to four sequence types (STs), including ST1076 (62.2%), ST463 (21.6%), ST646 (10.8%), and ST3393 (5.4%). To our knowledge, this is the first report of *bla*_KPC-3_ in *P. aeruginosa* ST463, ST646 and ST3393. All isolates exhibited extensive drug resistance, and 51.8% were resistant to ceftazidime-avibactam.

Phylogenetic analysis indicated that *bla*_KPC-3_ dissemination was driven by both clonal expansion and horizontal transmission. Comparative genomic analysis identified three kinds of *bla*_KPC-3_ -carrying plasmid. A transferable IncP-2 megaplasmid was widely distributed among ST1076, ST646, and ST3393 isolates, whereas non-transferable IncP-10 plasmids were primarily restricted to ST463. The genetic environments and plasmid backbones of *bla*_KPC-3_ were highly conserved and closely related to those of *bla*_KPC-2_ and its variants, suggesting evolution from pre-existing *bla*_KPC-2_-associated plasmids. Virulence analysis demonstrated marked heterogeneity across lineages.

ST463 isolates co-harbored *exoU* and *exoS*, exhibited enhanced biofilm formation and pyocyanin production, and caused significantly higher mortality in the *G. mellonella* infection model, indicating a hypervirulent phenotype.

**Conclusions:** The *bla*_KPC-3_ is becoming an increasingly important determinant of carbapenem resistance in *P. aeruginosa* in China. The IncP-2 megaplasmid and IncP-10 plasmid derived *bla*_KPC-3_ spread across multiple lineages. Continuous genomic surveillance and enhanced infection control measures are urgently needed to prevent its further prevalence in clinical settings.

## Introduction

*Pseudomonas aeruginosa* is a notorious opportunistic pathogen and a leading cause of healthcare-associated infections worldwide. It is frequently associated with severe clinical conditions, including pneumonia, bloodstream infections, and urinary tract infections, resulting in substantial morbidity and mortality[1]. Owing to its intrinsic resistance and remarkable genomic plasticity, the treatment of *P. aeruginosa* infections remains a major clinical challenge. Although carbapenems have long served as key therapeutic agents for managing multidrug-resistant *P. aeruginosa* infections, the global prevalence of carbapenem-resistant *P. aeruginosa* (CRPA) has markedly compromised their clinical effectiveness. Recent epidemiological data from the European Centre for Disease Prevention and Control (ECDC) showed that 16% of participating countries reported CRPA rates ≥50%, while the US CDC documented increasing hospital-onset multidrug-resistant *P. aeruginosa* infections after 2019[2, 3]. Consequently, the World Health Organization (WHO) has designated CRPA as a high-priority pathogen requiring urgent surveillance and therapeutic development[4].

Carbapenem resistance in *P. aeruginosa* arises through a combination of chromosomal and acquired mechanisms, including the inactivation of OprD outer membrane porin, overexpression of efflux pumps, hyperproduction of chromosomal AmpC β-lactamases, and the acquisition of various carbapenemase genes[1, 5].

Historically, carbapenemase-producing *P. aeruginosa* constituted a smaller proportion of CRPA isolates; however, this proportion has been steadily increasing over the past two decades[5]. Among these carbapenemases, metallo-β-lactamases (MBLs), such as VIM, IMP, and NDM, are the most prevalent in CRPA, with globally disseminated high-risk clones, such as ST235, ST111, and ST175, largely driving the epidemiological landscape[1, 6–8]. In contrast, *Klebsiella pneumoniae* carbapenemase (KPC), a class A serine β-lactamase widely disseminated in Enterobacterales, was previously considered uncommon in *P. aeruginosa*[5].

However, the epidemiology of KPC-producing *P. aeruginosa* has changed substantially over the past decade[9]. In China, *bla*_KPC-2_ has emerged as one of the most prevalent carbapenemase genes in CRPA and is frequently associated with the high-risk ST463 lineage[10]. Compared with *bla*_KPC-2_, *bla*_KPC-3_ differs by a single amino acid substitution (H272Y) but exhibits enhanced hydrolytic activity against carbapenems and oxyimino-cephalosporins, posing an even greater threat to clinical novel β-lactamase inhibitor combinations[11, 12]. Despite its clinical significance, *bla*_KPC-3_ remains rarely reported in CRPA worldwide, and its molecular epidemiology, transmission dynamics, and associated plasmid backgrounds are poorly understood. Recently, sporadic reports of *bla*_KPC-3_-producing *P. aeruginosa* have emerged in China, suggesting that this resistance determinant may be undergoing local expansion[13, 14]. In this study, we comprehensively analyzed the prevalence, molecular epidemiology, resistance and virulence characteristics of *bla*_KPC-3_ -producing CRPA isolates collected in China. We further explored the dissemination of *bla*_KPC-3_ across distinct clonal lineages and characterized the mobile genetic elements associated with its transmission.

## Materials and methods

### Strains and clinical data collection

During the period from January 2023 to January 2024, a total of 65 non-duplicate clinical CRPA isolates were collected from a tertiary teaching hospital in Zhejiang Province, China. All the isolates were subjected to species identification using MALDI-TOF MS technology (bioMérieux, Marcy l’Etoile, France). Demographic characteristics and clinical information of the corresponding patients were retrospectively retrieved from electronic medical records.

### Antimicrobial susceptibility testing

Minimum inhibitory concentrations (MICs) of 15 antimicrobial agents were determined using the broth microdilution method. Antimicrobial susceptibility results were interpreted according to the Clinical and Laboratory Standards Institute (CLSI) guidelines[15]. *P. aeruginosa* ATCC 27853 served as quality control strain. The antimicrobial agents tested included cefazolin (CFZ), Cefuroxime (CXM), ceftazidime (CAZ), cefepime (FEP), meropenem (MEM), imipenem (IPM), aztreonam (ATM), Tobramycin (TOB), amikacin (AMK), ciprofloxacin (CIP), levofloxacin (LEV), piperacillin-tazobactam (TZP), cefoperazone-sulbactam (SCF), amoxicillin/clavulanic (AMC), and ceftazidime-avibactam (CZA).

### Whole-genome sequencing and bioinformatics analysis

Genomic DNA was extracted using a genomic DNA isolation kit (Tiangen, DP305, China). Short-read whole-genome sequencing was performed on the Illumina NovaSeq 6000 platform (Illumina Inc., San Diego, CA, USA) to generate 150 bp paired end reads. Raw data were quality-filtered using Trimmomatic v0.39, and *de novo* assembled with SPAdes v3.15.2[16, 17]. To obtain complete genome sequences, four representative isolates were further subjected to long-read sequencing using the Oxford Nanopore MinION platform (Oxford Nanopore Technologies, Oxford, UK). Hybrid genome assembly was performed by combining Illumina short reads and Nanopore long reads using Unicycler v0.4.8[18].

Antimicrobial resistance genes were identified using ResFinder 4.1 and the CARD database. Virulence genes were identified based on VFDB database. Multilocus sequence typing (MLST) was performed using the MLST 2.1 servers available at the Center for Genomic Epidemiology. A phylogenetic tree based on single nucleotide polymorphisms (SNPs) was constructed with Snippy-multi (https://github.com/tseemann/snippy) and Fasttree[19].

### Conjugation experiments

Conjugation assays were performed to assess plasmid transferability. *P. aeruginosa* PAO1^AprR^ and *E. coli* EC600 were used as the recipients, and the *bla*_KPC-3_-carrying isolates served as donors. Transconjugants were selected on Mueller-Hinton agar plates which containing apramycin (100 μg/mL) or rifampicin (600 μg/mL), supplemented with meropenem (2 μg/mL). The transconjugants were identified by MALDI-TOF-MS, and the presence of the *bla*_KPC-3_ gene in transconjugants was confirmed by PCR and Sanger sequencing.

### Biofilm formation assays

Biofilm formation was semi-quantified using the crystal violet staining assay. Briefly, overnight bacterial cultures were diluted with Luria-Bertani (LB) broth and incubated in 96-well microtiter plate for 24 h at 37°C. Subsequently, microwells were gently washed three times with phosphate-buffered saline (PBS) to remove non-adherent cells. The attached biofilm was fixed with methanol and stained with 1% crystal violet. Plates were rinsed with running water to remove excess stain and air-dried. Then, 100 μL of 30% acetic acid were added to each well to solubilize the bound crystal violet, and biofilm biomass was quantified by measuring the optical density (OD) at 600 nm. Sterile LB broth was used for the negative control. Experiments were conducted in triplicates and independently repeated three times.

### Pyocyanin assay

*P. aeruginosa* isolates were cultured in LB broth at 37^◦^C with 220 rpm shaking for 16 h. Bacterial cultures were centrifuged at 12,000 × g, and the supernatant was collected for pyocycin extraction. Briefly, supernatants were extracted twice with chloroform (5:3, v/v) until a greenish-blue organic phase formed. After centrifugation (10,000 × g for 10 min), the chloroform phase was transferred to a new tube containing 0.2 M HCl and vortexed until the solution turned pink. The upper aqueous phase (pink) was then collected, and pyocyanin production was quantified by measuring absorbance at 520 nm.

### *Galleria mellonella* infection model

The virulence of *P. aeruginosa* isolates was assessed using the *G. mellonella* infection model. Bacterial cells in the logarithmic growth phase were harvested and adjusted to approximately 1 × 10^5^ CFU/ml in sterile PBS. Each larva was inoculated with 10 μl of bacterial suspension via the last left proleg, with 20 larves included in each experimental group. After infection, larvae were incubated at 37°C and monitored for survival over 48 h. Sterile PBS was used as the negative control, while *P. aeruginosa* PAO1 strain was used as the moderate virulence control.

## Results

### High prevalence of *bla*_KPC-3_ in clinical CRPA isolates

Among the 65 CRPA isolates, genomic analysis identified 37 isolates (56.9%) carrying carbapenemase gene *bla*_KPC-3_. MLST analysis assigned these *bla*_KPC-3_-positive isolates to four ST lineages, including ST1076 (62.2%, 23/37), ST463 (21.6%, 8/37), ST646 (10.8%, 4/37), and ST3393 (5.4%, 2/37) **(Figure 1).**

**Figure 1.**
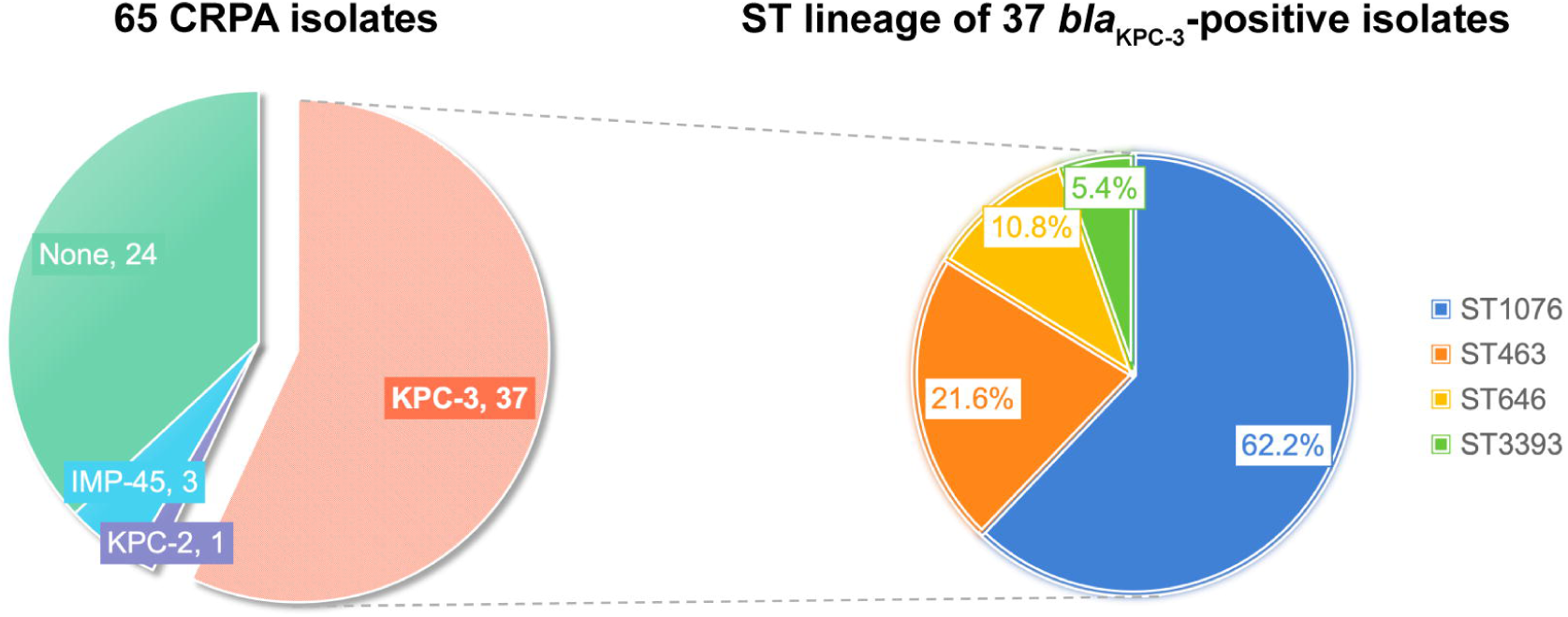
Prevalence and clonal distribution of *bla*_KPC-3_-positive CRPA clinical isolates. The left pie chart illustrates the distribution of carbapenemase genes among the 65 clinical CRPA isolates in this study. The right multilayer pie chart shows the multilocus sequence typing (MLST) distribution of the 37 *bla*_KPC-3_-positive strains.

Although ST1076 was the predominant lineage, the identification of *bla*_KPC-3_ in phylogenetically distant STs suggests that its dissemination is not solely driven by clonal expansion and may involve horizontal gene transfer.

These isolates were primarily recovered from respiratory tract (67.6%, 25/37) and bloodstream samples (13.5%, 5/37). Most patients were elderly males (67.6%; median age, 72 years) with severe underlying diseases, ICU admission, and invasive procedures. Broad-spectrum β-lactams were frequently administered prior or during infection, including piperacillin/tazobactam (n=20), cefoperazone/sulbactam (n=15), meropenem (n=12), and ceftazidime/avibactam (n= 4). Clinical outcomes were poor, with only nine patients (24.3%) showing improvement, while thirteen patients (35.1%) died during hospitalization (Table S1).

### Antimicrobial susceptibility and resistance profiles

Due to strain availability, a subset of 27 isolates was ultimately included in subsequent analyses. Antimicrobial susceptibility testing showed that all *bla*_KPC-3_-harboring isolates exhibited a multidrug-resistant phenotype, with extensive resistance to cephalosporins, carbapenems, aztreonam and β-lactamase inhibitor combinations **(Figure 2)**. Fluoroquinolone resistance was common, with 66.7% and 70.3% of isolates resistant to ciprofloxacin and levofloxacin, respectively, which largely driven by ST463 (100% resistance) and ST1076 (87.5% resistance). Notably, resistance to ceftazidime-avibactam (CZA) was observed in 51.8% of isolates, with all ST463 isolates exhibiting high-level resistance (MIC ≥32 μg/ml). In contrast, aminoglycosides remained relatively effective. No isolates exhibited resistance to amikacin, and only two isolates were resistant to tobramycin **(Figure 2, Table S2)**.

**Figure 2.**
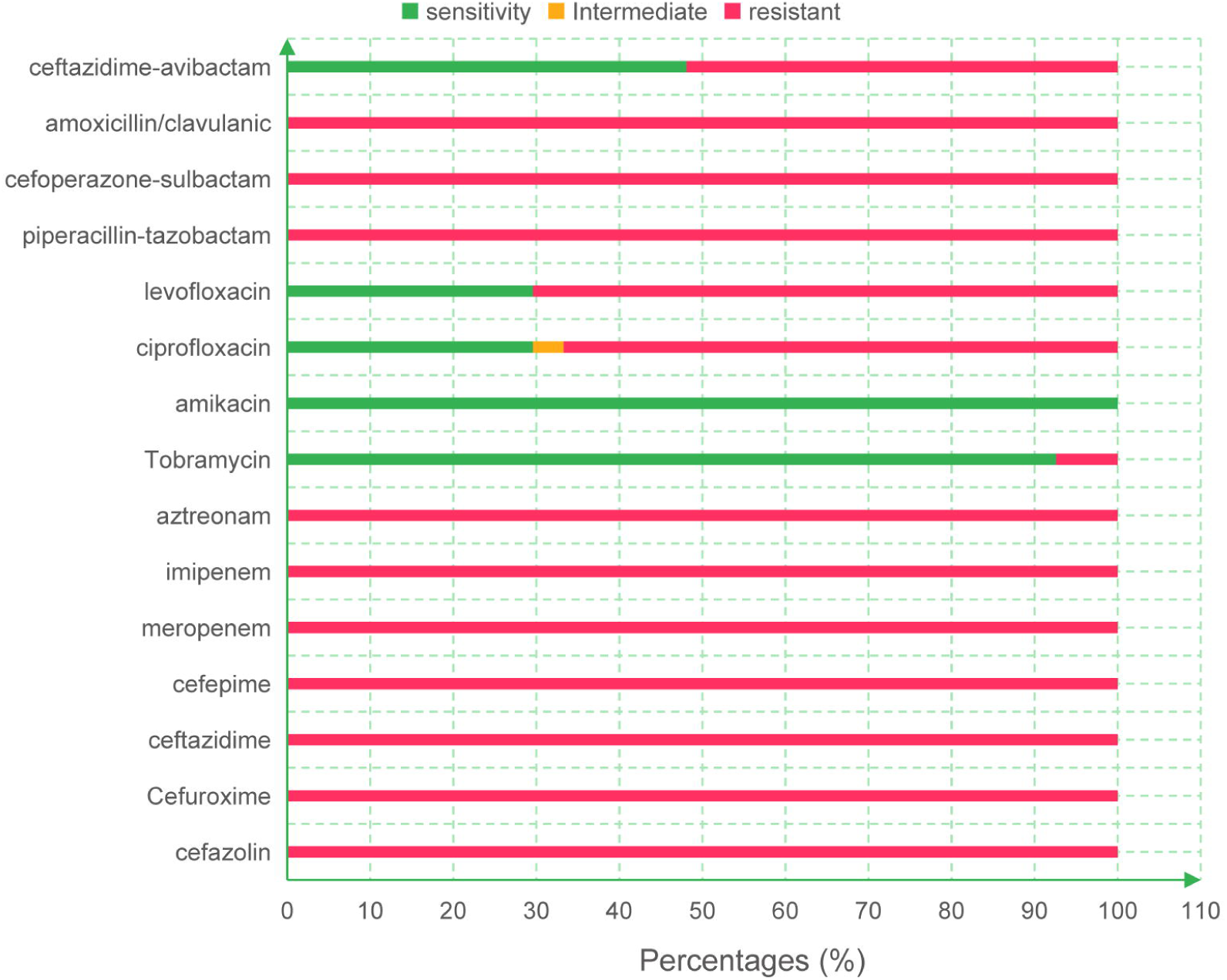
Antimicrobial susceptibility of 27 *bla*_KPC-3_-positive CRPA isolates.

### Antimicrobial resistance and virulence determinants

As shown in **Figure 3B**, all isolates possessed a core set of intrinsic chromosomal resistance determinants, including the aminoglycoside modifying enzyme *aph* (3′)-IIb, the fosfomycin resistance gene *fosA*, the chloramphenicol acetyltransferase *catB7*, and several multidrug efflux pumps (MexAB-OprM, MexCD-OprJ, and MexEF-OprN).

**Figure 3.**
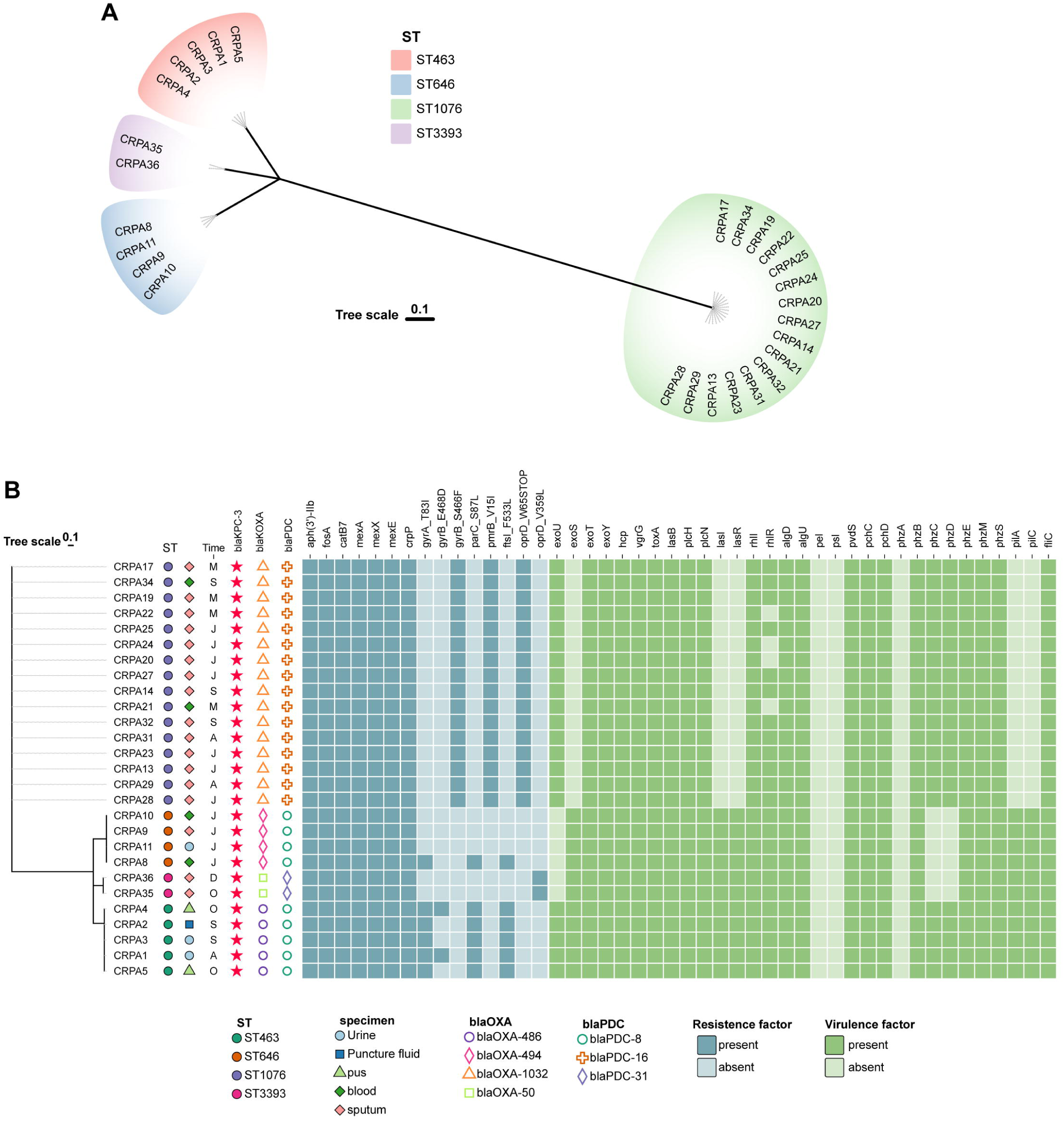
Phylogenetic analysis and molecular characteristics of KPC-3 producing *P. aeruginosa* isolates. A maximum-likelihood phylogenetic tree was constructed based on core-genome SNPs. The presence or absence of resistance gene and virulence genes are indicated by filled and empty squares, respectively.

**Figure 4.**
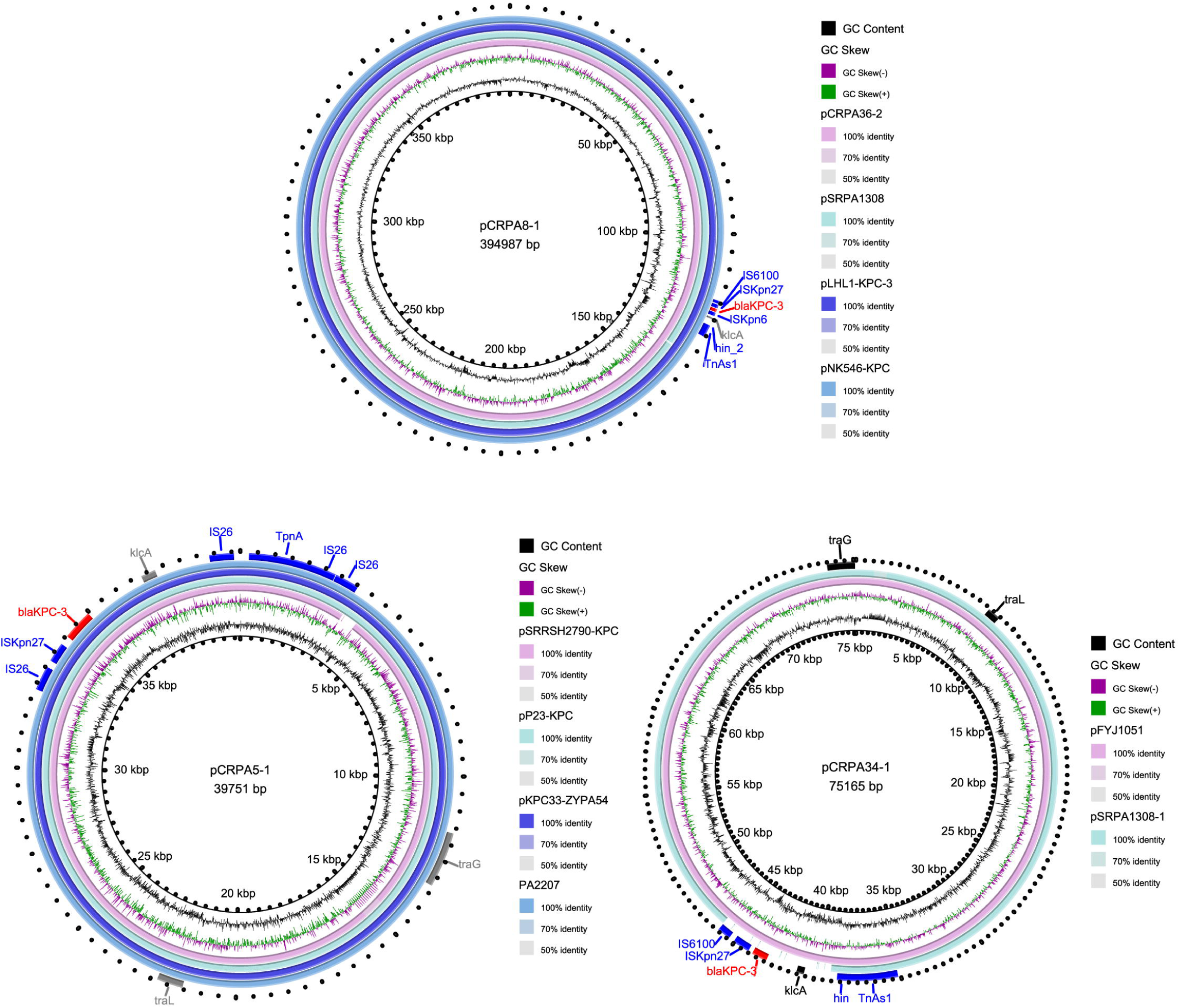
Comparative analysis of the *bla*_KPC-3_-haboring plasmids in this study and other similar plasmids available from the NCBI databases. (A). pCRPA8-1 was used as the reference plasmid to perform the genome alignment with pCRPA36-2, pSRPA1308, pLHL1-KPC-3 and pNK546-KPC. (B). pCRPA5-1 was used as the reference plasmid to perform the genome alignment with pSRRSH2790-KPC, pP23-KPC, and pKPC33-ZYPA54. (C). pCRPA34-1 was used as the reference plasmid to perform the genome alignment with pFYJ1051 and pSRPA1308-1. The concentric circles from the inside to the outside represent each plasmid, as shown in the right column. Colored regions demonstrate homologous sequences, whereas gaps represent regions lacking sequence similarity.

**Figure 5.**
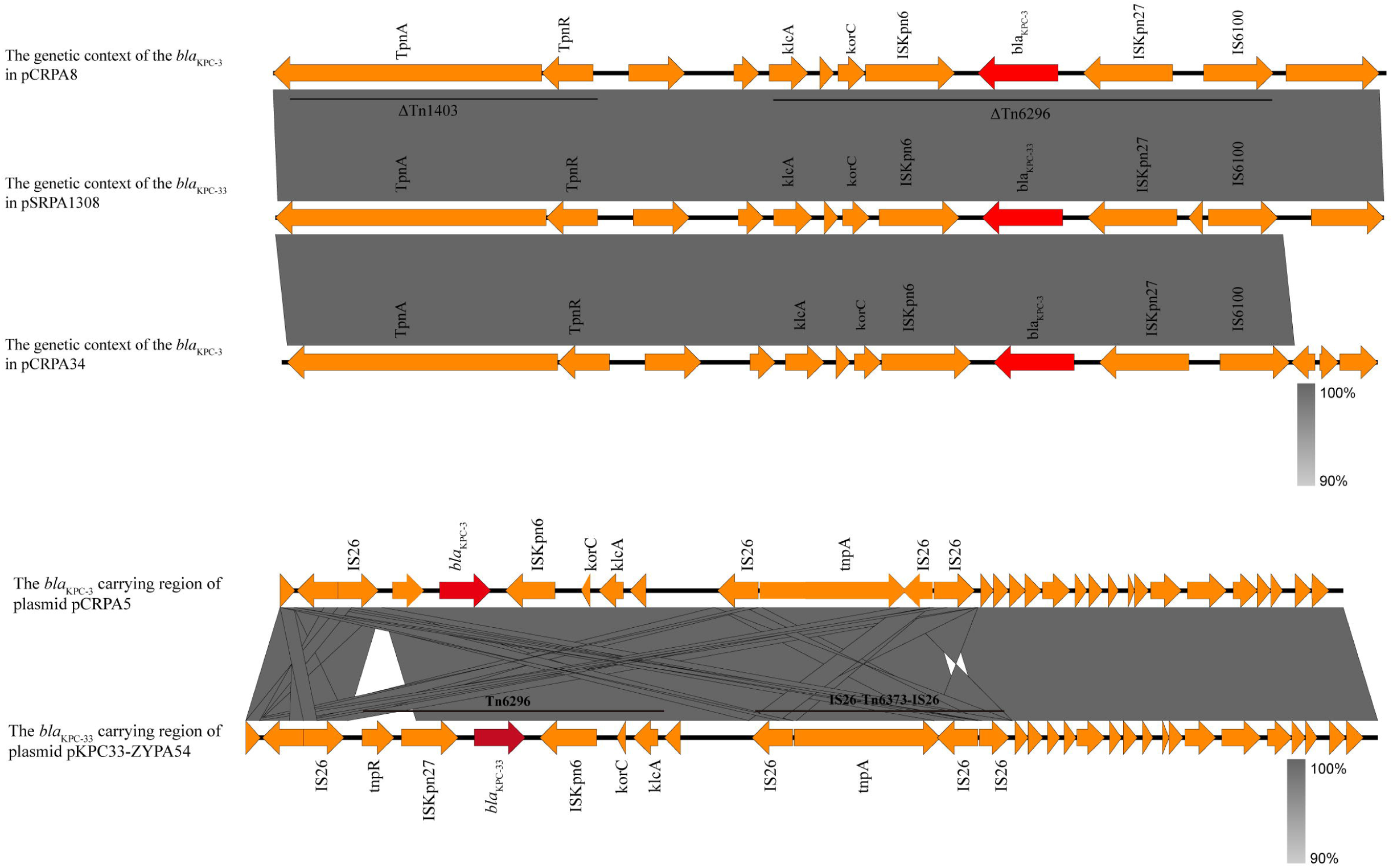
The genetic context of the *bla*_KPC-3_ gene on the plasmid. Arrows represent complete or truncated genes, with arrowheads indicating the direction of transcription. Gray shading indicates homologies region shared among corresponding genetic loci.

**Figure 6.**
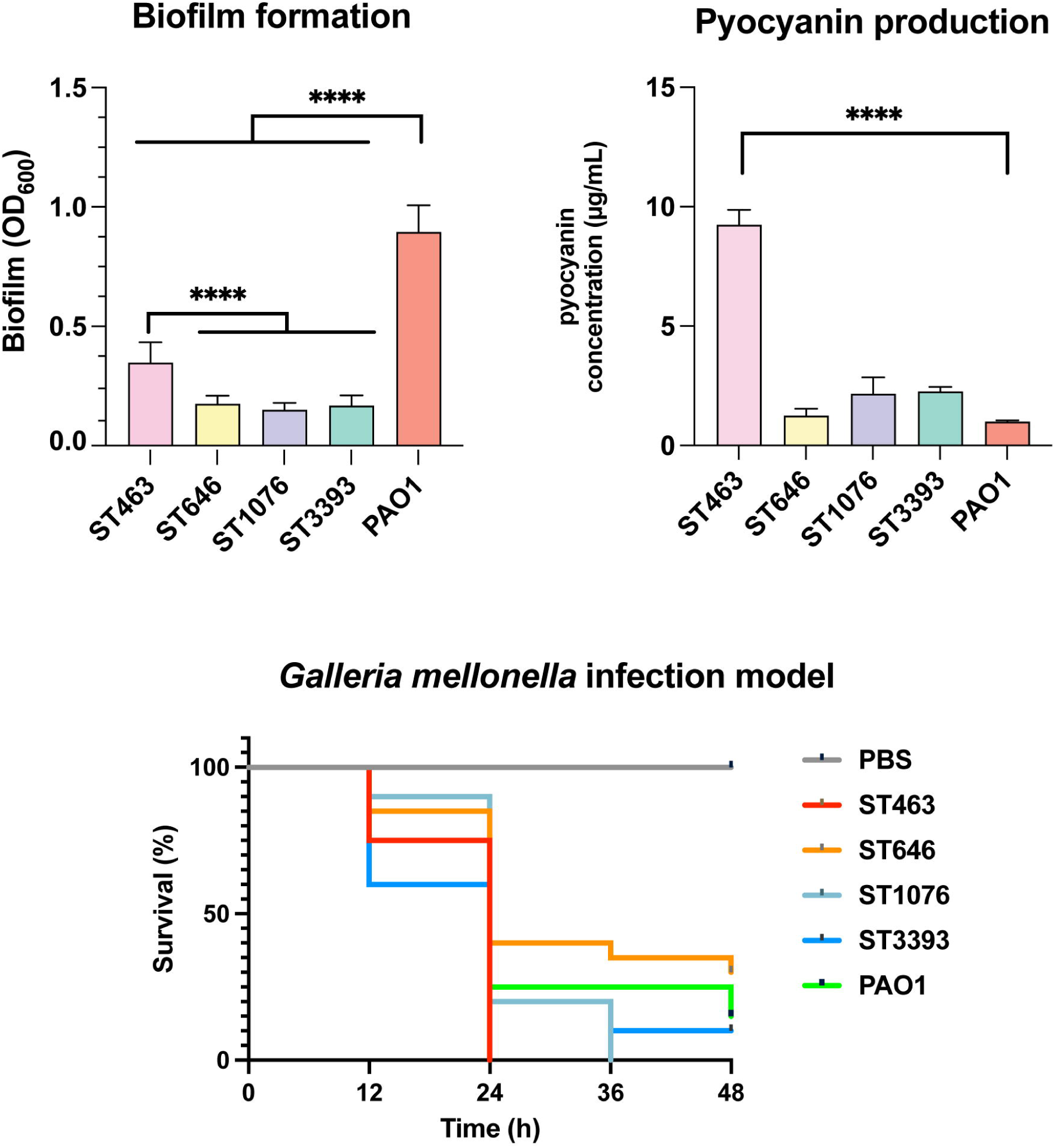
*In vitro* and *in vivo* virulence assessment of *bla*_KPC-3_-producing *P. aeruginosa* isolates from different STs. **(A)** Quantification of biofilm formation by crystal violet staining (n=27). **(B)** Measurement of pyocyanin production (n=27). **(C)** Survival analysis of the *G. mellonella* infection model challenged with one representative isolate from each ST. PAO1 as the medium virulence control. Each infection assay was performed independently three times.

Additionally, the intrinsic β-lactamase genes *bla*_PDC_ and *bla*_OXA_ were identified in all genomes, with their allelic variants exhibiting high lineage specificity. Specifically, *bla*_OXA-50_, *bla*_OXA-1032_, *bla*_OXA-486_, and *bla*_OXA-494_ were restricted to ST3393, ST1076, ST463 and ST646 isolates, respectively. Similarly, *bla*_PDC_-_8_ was detected in ST463 and ST646, *bla*_PDC_-_16_ in ST1076, and *bla*_PDC_-_31_ in ST3393 isolates. All isolates carried the ciprofloxacin-modifying enzyme gene *crpP*, while quinolone resistance-associated mutations (*gyrB_*S466F, *gyrB*_E468D, *gyrA*_T83I, and *parC*_S87L) were mainly detected in ST463 and ST1076, consistent with their elevated fluoroquinolone resistance rates. Notably, all ST1076 isolates harbored *pmrB*_V15I, a mutation associated with altered colistin susceptibility. In addition, all ST463 isolates harbored *ftsI*_F533L, whereas alterations in *oprD* were primarily found in ST1076 (*oprD*_W65STOP) and ST3393 (*oprD*_V359L), potentially synergizing with *bla*_KPC-3_ to confer enhanced β-lactam and carbapenem resistance[20].

Virulence profiling revealed a conserved repertoire of key virulence determinants, including type VI secretion system (T6SS) components *hcp* and *vgrG*, toxin/enzyme/protease-related genes (*toxA*, *lasB*, *plcH*, *plcN*), antiphagocytic operons (*algD*, *algU*), iron acquisition regulators (*pvdS, pchC/D*), phenazines biosynthesis clusters (*phzB/E/M/S*), and the flagellin gene *fliC*. The type III secretion system (T3SS) represents a major determinant of acute virulence in *P. aeruginosa*, with ExoU and ExoS as its key effectors. The type III secretion system *exoU* and *exoS* were detected in 77.8% (21/27), and 40.7% (11/27) of isolates, respectively. ST646 and ST3393 isolates were *exoU−/exoS+*, ST1076 isolates were *exoU+/exoS−*. Notably, ST463 isolates simultaneously harbored *exoU* and *exoS*, indicating their hypervirulence.

### Phylogenetic analysis

To evaluate the genetic relatedness of the CRPA isolates, we performed core-genome SNP analysis and constructed a phylogenetic tree based on SNP differences.

Phylogenetic analysis demonstrated that each ST formed a distinct, well-defined phylogenetic clade, with isolates of the same ST clustering tightly and exhibiting high internal genomic homology **(Figure 3A)**. Pairwise SNP analysis further elucidated the fine-scale genetic variation with each lineage. Based on a conservative threshold where isolates differing by fewer than 25 SNPs were considered to belong to the same transmission cluster, our data strongly supported clonal transmission within the hospital[21]. Overall, intra-ST pairwise distances ranging from 0-103 SNPs **(Figure S1)**. Specifically, the ST646 and ST3393 isolates exhibiting minimal genetic divergence, differing by only 0-6 SNPs. The predominant ST1076 lineage showed pairwise SNP differences ranging from 1-70 (median, 30 SNPs), consistent with recent clonal expansion and ongoing nosocomial circulation. In contrast, the ST463 lineage displayed a boarder range of genetic variation, with SNP differences spanning from 41 to 103. This higher diversity suggests a longer evolutionary history or multiple introduction events within the hospital environment.

### Characterization of *bla*_KPC-3_–harboring plasmids

Analysis of Illumina short-read sequencing data revealed that *bla*_KPC-3_ was situated on three kinds of plasmids among the 27 clinical isolates. Notably, an IncP-2 pSRPA1308-like plasmid emerged as the predominant *bla*_KPC-3_ carrying platform, exhibiting broad distribution across ST646, ST3393, and ST1076 isolates. An exception was observed in one ST1076 isolate (CRPA34), in which *bla*_KPC-3_ was located on an IncP-10 pSRPA1308-1-like plasmid **(Table S3)**. In contrast, all ST463 isolates carry *bla*_KPC-3_ on an IncP-10 pSRRSH2790-KPC-like plasmid. To further characterize the genomic architecture, we subsequently conducted long-read genome sequencing for four representative strains (CRPA5, CRPA8, CRPA34 and CRPA36). All four isolates were found to possess a ∼7 Mb circular chromosome and 1-4 plasmids of varying sizes **(Table S3)**. Their *bla*_KPC-3_ gene is plasmid-borne, and represented the sole antimicrobial resistance gene on each plasmid.

Consistent with the short-read sequencing analysis, long-read sequencing confirmed that pCRPA8-1 and pCRPA36-2 were identical. Both plasmids were 394,987 bp in size, contained 434 predicted open reading frames, and had a G+C content of 56.57%, belonging to an IncP-2 megaplasmid family. Therefore, we chose pCRPA8-1 as the representative plasmid for further analysis. Comparative plasmid analysis showed that the pCRPA8-1 was almost identical to plasmid pSRPA1308 (GenBank: CP158571) carrying *bla*_KPC-33_ from ST463, plasmid pLHL1-KPC-3 (GenBank: CP099961), the first reported IncP-2 *bla*_KPC-3_-carrying plasmid from clinical *P. aeruginosa*, and pNK546-KPC (GenBank: MN433457) carrying *bla*_KPC-2_ from *P. aeruginosa* isolate, with 100% query coverage and 100% nucleotide identity.

The IncP-10 plasmid pCRPA5-1 and pCRPA34-1 were relatively small (39,751 and 75,165 bp in size, respectively), with similar G+C contents, and harboring 46 and 82 predicted open reading frames, respectively. BLASTn analysis revealed that pCRPA5-1 shared high sequence similarity with multiple plasmids carrying *bla*_KPC-2_, *bla*_KPC-33_ and *bla*_KPC-90_, such as pSRRSH2790-KPC (GenBank: CP077995), pP23-KPC (GenBank: CP065418), pKPC33-ZYPA54 (GenBank: OK105106) and PA2207 (GenBank: CP080290), showing 93% to 100% query coverage and nucleotide identity. Information on the 24 related plasmids is summarized in **Table S4**. Similarly, pCRPA34-1 exhibited high sequence similarity to pFYJ1051 (GenBank: PP712856) and pSRPA1308-1 (GenBank: CP158572), which carry *bla*_KPC-2_ and *bla*_KPC-33_ respectively.

These traits suggest that the *bla*_KPC-3_-bearing plasmids in this study share nearly identical backbones with plasmids of *bla*_KPC-2_ and its variants previously associated with carbapenem and CZA resistance of *P. aeruginosa* in China, indicating that these *bla*_KPC-3_-bearing plasmids potentially evolved independently from established *bla*_KPC-2_-carring plasmids before becoming epidemic within the local region.

Genetic environment analysis indicated two conserved genetic platforms surrounding the *bla*_KPC-3_ among our isolates. The first platform showed identified genetic contexts flanking *bla*_KPC-3_ in the IncP-2 megaplasmid pCAPA8-1 and the IncP-10 plasmid pCAPA34-1. In both plasmids, the *bla*_KPC-3_ gene is embedded within a conserved IS*6100*-ΔTn*6296*-ΔTn*1403* structure. This structural identity suggested that this *bla*_KPC-3_-bearing mobile genetic element may be capable of inter-plasmid transfer. The second platform was identified in IncP-10 plasmid pCRPA5-1, where the *bla*_KPC-3_ resistance gene is located in a genetic structure of IS6-IS*Kpn27*-*bla*_KPC-3_-IS*26*- *korC-klcA*-IS*26*-*TnpA*-IS*26*-IS*26*. This region contains two IS*26* units, including IS*26-bla*_KPC_-IS*26* and IS*26*-Tn*6373*-IS*26*. A similar genetic organization had previously been reported in plasmids of ST463 CRPA isolates in China, with pKPC33-ZYPA54 as a representative example. Interestingly, sequence alignment showed that pCRPA5-1 harbored a truncated Tn*6376* element lacking transposase gene *tnpA*, likely mediated by IS*26* insertion. In addition, a large-scale inversion event encompassing the backbone genes (*korC* and *klcA*) and IS*26*-Tn*6376*-IS*26* module was identified, indicating a localized genomic rearrangement within this plasmid.

The horizontal transfer potential of the *bla*_KPC-3_ -carrying plasmids was assessed through conjugation experiments using apramycin-resistant *P. aeruginosa* PAO1^AprR^ and rifampicin-resistant *E. coli* EC600 as the recipients. None of the *bla*_KPC-3_ -carrying plasmid could be transferred to *E. coli* EC600, whereas the IncP-2 megaplasmids pCRPA8-1 and pCRPA36-2 were successfully conjugated into PAO1^AprR^, with conjugation frequencies ranging from 3.6×10^-5^ to 7.0×10^-6^, confirming the transferability of these IncP-2 megaplasmids with *P. aeruginosa* species. By comparison, the IncP-10 plasmids pCRPA5-1 and pCRPA34-1 failed to conjugate to PAO1^AprR^ after three independent attempts. Compared with the recipient PAO1^AprR^, transconjugants PAO1^AprR^/pCRPA8-1 and PAO1^AprR^/pCRPA8-1 exhibited increased resistance to cefepime, ceftazidime, aztreonam, meropenem and imipenem to varying degrees. Notably, the MICs of CZA and cefiderocol increased approximately fourfold in both transconjugants, indicating that these *bla*_KPC-3_-carrying IncP-2 megaplasmids can reduce susceptibility to both agents in *P. aeruginosa*.

### Virulence assessment *in vitro* and *in vivo*

To characterize the pathogenicity of the *bla*_KPC-3_-positive CRPA, we assessed their biofilm formation, pyocyanin secretion, and in vivo lethality using a *G. mellonella* infection model. In the *in vitro* phenotypic assays, ST463 isolates demonstrated a prominent hypervirulent profile. Specifically, ST463 yielded the highest biofilm biomass (0.35 ± 0.08), which was significantly higher than that of the ST646 (0.18 ± 0.03), ST1076 (0.15 ± 0.03), and ST3393 (0.17 ± 0.04) isolates (*p* < 0.05), although all of these STs isolates exhibited lower biofilm capacity than the reference strain PAO1. Consistently, ST463 isolates exhibited a marked elevation in pyocyanin production compared with the other STs (*p* < 0.05). These findings were well corroborated by *G. mellonella* survival assays. At 24 h post-infection, ST463 caused 100% larval mortality, compared with 75% mortality for the moderate virulence control PAO1, indicating significantly enhanced virulence (*p* < 0.0001). Meanwhile, ST1076 and ST3393 exhibited medium virulence, producing mortality rates comparable to that of PAO1. In contrast, ST646 showed the lowest virulence among all tested lineages, with only 60% larval mortality at 24 h and 65% mortality at 48 h. Collectively, these results indicate substantial heterogeneity in virulence among *bla*_KPC-3_-producing CRPA lineages, with ST463 representing the most hypervirulent clone and ST646 displaying a relatively hypovirulent phenotype.

## Discussion

The present study revealed an unexpectedly regional prevalence of *bla*_KPC-3_ among clinical CRPA isolates in eastern China, where it was detected in 56.9% of isolates. Although *bla*_KPC-2_ remains the predominant KPC variant in *P. aeruginosa* globally, *bla*_KPC-3_ has only rarely been reported in this pathogen, making its high prevalence in this study particularly noteworthy[9, 10, 22]. The earliest report of *bla*_KPC-3_ -producing *P. aeruginosa* in China described five ST1076 isolates recovered from two patients in Hangzhou in 2021[14]. Consistent with their findings, ST1076 was the predominant of *bla*_KPC-3_-carrying lineage in our collection. Additionally, we also identified *bla*_KPC-3_ in ST463, ST646 and ST3393, representing, to our knowledge, the first report of *bla*_KPC-3_ in these lineages. The dissemination of *bla*_KPC-3_ across multiple lineages suggests that it has become established in the local CRPA population and may represent an emerging epidemiological threat in China.

The emergence and dissemination of *bla*_KPC-3_ may reflect adaptive evolution under antimicrobial selection pressure. Compared with KPC-2, KPC-3 has been reported to exhibit enhanced hydrolytic activity against several β-lactams, including carbapenems and extended-spectrum cephalosporins[11]. CZA and cefiderocol are important therapeutic options for multidrug-resistant *P. aeruginosa*, and exposure to these agents could favor variants with reduced susceptibility[23–25]. In our study, introduction of the *bla*_KPC-3_-carrying IncP-2 megaplasmids into PAO1 increased the MICs of both CZA and cefiderocol by approximately 4-fold, suggesting that these plasmids can directly reduce susceptibility to these clinically important agents. Moreover, comparative genomic analysis demonstrated that all *bla*_KPC-3_-carrying plasmids identified in this study shared highly conserved backbones and genetic environments with previously reported *bla*_KPC-2_ and its variant-bearing plasmids[6, 14, 26, 27].

These findings support the possibility that *bla*_KPC-3_ evolved from pre-existing *bla*_KPC-2_-associated mobile genetic elements through point mutation, followed by local dissemination. Given the widespread use of carbapenems and the increasing clinical application of newer antipseudomonal agents, the ability of *bla*_KPC-3_-carrying plasmids to reduce susceptibility to both carbapenems and these agents may provide an adaptive advantage in certain clinical settings[28, 29].

All *bla*_KPC-3_-positive isolates exhibited extensively drug-resistant phenotypes, and 51.8% were resistant to CZA, a rate substantially higher rate than that reported for KPC-2-producing *P. aeruginosa[30]*. Transfer of the *bla*_KPC-3_-carrying IncP-2 plasmids into PAO1 increased the CZA MIC by approximately fourfold, demonstrating a direct contribution to reduced CZA susceptibility. Consistent with this finding, the CZA MICs of most isolates were already at or above the susceptibility breakpoint (8/4 mg/L). However, the higher MICs observed in clinical isolates indicate that additional chromosomal mechanisms are likely required[31]. As anticipated, multiple intrinsic efflux systems were present in the isolates, and alterations in *oprD*, a major determinant of carbapenem susceptibility, were also detected. Notably, no typical “seesaw effect” was observed. Unlike some KPC variants, such as KPC-33, KPC-71 or KPC-78[32], for which increased CZA resistance may be accompanied by increased carbapenem susceptibility in *P. aeruginosa*, the CZA-resistant isolates in our study remained highly resistant to carbapenems. The coexistence of these chromosomal determinants with *bla*_KPC-3_ may compensate for any fitness cost associated with altered carbapenemase activity and maintain high-level carbapenem resistance[33], thereby substantially limiting available therapeutic options.

The distribution of *bla*_KPC-3_ across genetically distinct lineages suggested a role for horizontal gene transfer in its dissemination. While SNP analysis supported clonal expansion of ST646, and ST3393 within the hospital, the detection of *bla*_KPC-3_ in genetically distinct ST463 and ST1076 isolates cannot be readily explained by vertical transmission alone. Plasmid analysis revealed two major *bla*_KPC-3_-carrying platforms: IncP-2 megaplasmids and IncP-10 plasmids[34]. The IncP-2 plasmids were widely distributed across multiple sequence types, displayed highly conserved genomic structures, and were successfully transferred to *P. aeruginosa* PAO1 in conjugation experiments, supporting their role as important vehicles for the horizontal dissemination of *bla*_KPC-3_. In contrast, IncP-10 plasmids were primarily confined to ST463 isolates and remained non-transferable under experimental conditions, suggesting a more restricted role in plasmid-mediated spread. The conserved genetic structures surrounding *bla*_KPC-3_, including IS26-, ISKpn27-, and ISKpn6-associated elements were closely related to previously described *bla*_KPC_-bearing mobile platforms[35], further supports the involvement of mobile genetic elements in its evolution and dissemination. Taken together, these findings suggest that *bla*_KPC-3_ dissemination has involved both plasmid-mediated horizontal transfer and subsequent adaptation within distinct bacterial backgrounds. The demonstrated conjugative capacity and broad distribution of the IncP-2 megaplasmids highlight the need to monitor transferable plasmids in addition to high-risk CRPA clones.

The acquisition of *bla*_KPC-3_ by ST463 is particularly concerning because this lineage is recognized as one of most successful high-risk CRPA clone in China[36, 37].

Previous studies have linked the expansion of ST463 to *bla*_KPC-2_-carrying plasmids and have demonstrated its enhanced virulence potential[37–39]. Consistent with these reports, all ST463 isolates in our study carried both *exoU* and *exoS* and exhibited enhanced biofilm formation, increased pyocyanin production, and significantly greater virulence in the *G. mellonella* infection model. The convergence of extensive antimicrobial resistance and hypervirulence within this lineage raises substantial clinical concern, as such strains may possess enhanced capacity for persistence, transmission, and severe infection[38, 39]. The identification of *bla*_KPC-3_ within this high-risk genetic background therefore represents a particularly alarming development.

Virulence phenotypes varied among the other *bla*_KPC-3_-producing lineages. ST1076 and ST646 showed moderate virulence, whereas ST3393 exhibited relatively low virulence in the *G. mellonella* model despite carrying multiple virulence-associated genes. These differences indicate that the phenotypic consequences of *bla*_KPC-3_ acquisition are strongly influenced by the genetic background of the host strain. The identification of *bla*_KPC-3_ in ST463, ST646 and ST3393 also expands the known clonal diversity of KPC-3-producing *P. aeruginosa* and highlights the potential for this resistance determinant to establish in lineages with different virulence characteristics.

In conclusion, this study identifies a noteworthy regional prevalence of *bla*_KPC-3_ among clinical CRPA isolates in eastern China and demonstrates its dissemination across multiple sequence types. The combination of a transferable IncP-2 megaplasmid, extensive antimicrobial resistance, and the hypervirulent ST463 lineage provides multiple routes for the persistence and spread of *bla*_KPC-3_. Continuous genomic surveillance of high-risk clones and transferable resistance plasmids is therefore essential to limit the further dissemination of *bla*_KPC-3_ in clinical settings.

---

**Figure S1 Pairwise SNP distance matrix of isolates within each ST (ST1076, ST463 and ST646).**

## CRediT authorship contribution statement

**Shanshan Wang**: Methodology, Investigation, Writing-original draft. **Meilan Li**: Investigation, Methodology. **Zhixuan Chen**: Investigation, Data curation. **Lan Chen**: Resources. **Xingbei Weng**: Resources, Funding acquisition. **Liang Chen**: Conceptualization, Software, Visualization, Writing-review & editing. **Bingjie Wang**: Conceptualization, Supervision, Writing-original draft, Writing-review & editing.

## Funding

This work was supported by the Ningbo Major Research and Development Plan Project (2024Z216).

## Data availability

The datasets underpinning the results of this research can be obtained from the corresponding author upon a justified request.

## Notes

### Competing Interest Statement

The authors have declared no competing interest.

