## Supplementary figures and images for "High prevalence of KPC-3 in carbapenem-resistant *Pseudomonas aeruginosa* across multiple clonal lineages in China"

### Figure S1

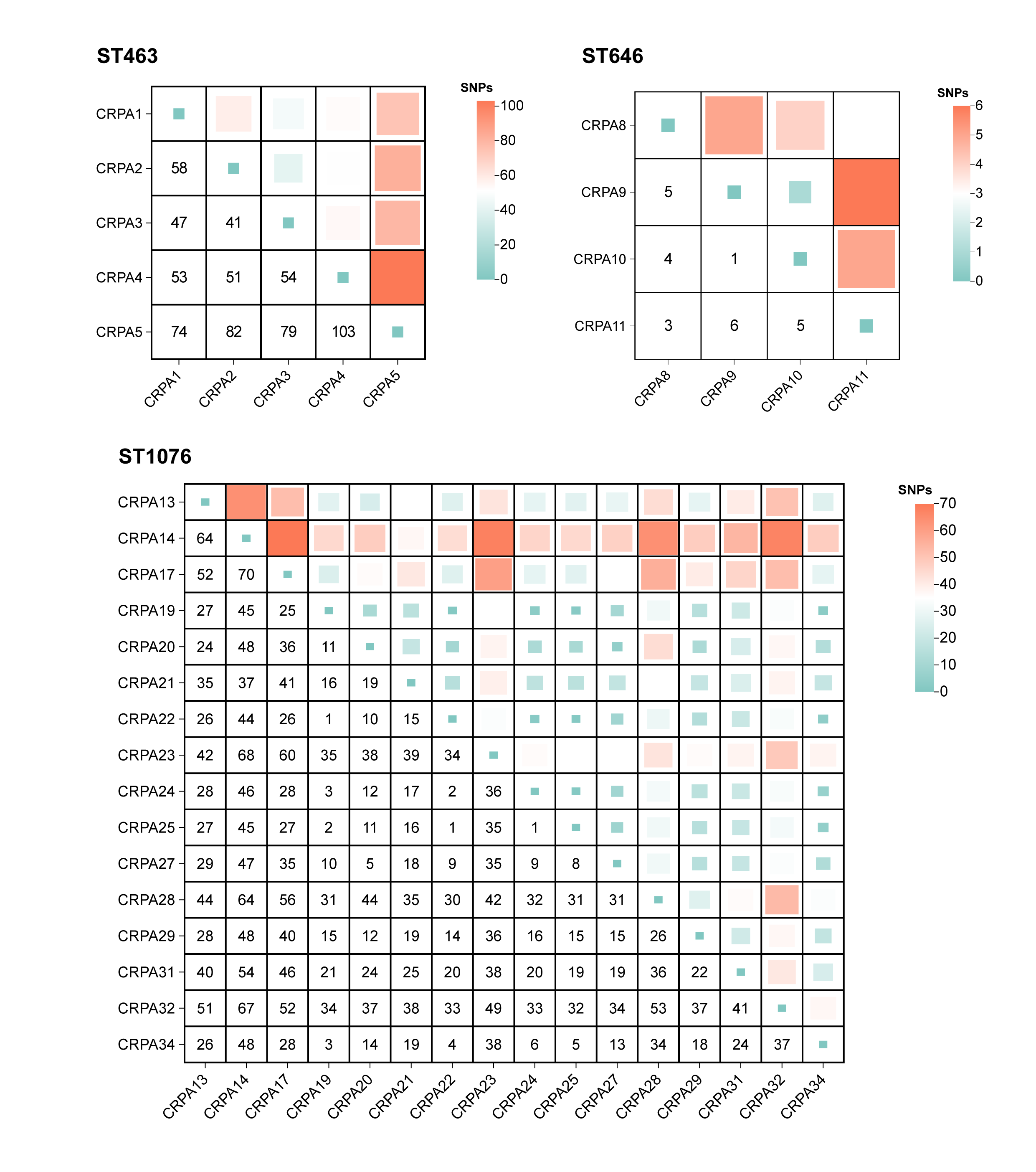
